# Electrophysiological indices of hierarchical speech processing differentially reflect the comprehension of speech in noise

**DOI:** 10.1101/2023.03.30.534927

**Authors:** Shyanthony R. Synigal, Andrew J. Anderson, Edmund C. Lalor

## Abstract

The past few years have seen an increase in the use of encoding models to explain neural responses to natural speech. The goal of these models is to characterize how the human brain converts acoustic energy into distinct linguistic representations that enable everyday speech comprehension. For example, researchers have shown that electroencephalography (EEG) data can be modeled in terms of acoustic features of speech, such as its amplitude envelope or spectrogram, linguistic features such as phonemes and phoneme probability, and higher-level linguistic features like context-based word predictability. However, it is unclear how reliably EEG indices of these speech feature representations reflect comprehension in different listening conditions. To address this, we recorded EEG from neurotypical adults who listened to segments of an audiobook in various levels of background noise. We modeled how their EEG responses reflected a range of acoustic and linguistic speech features and how this tracking varied with behavior across noise levels. EEG tracking of nearly all examined features showed SNR-dependent changes in unique variance explained, with the largest changes occurring for linguistic features. We hypothesized that only higher-level feature tracking would predict behavior but instead found that both high and low-level features were associated with behavioral scores depending on the noise level. EEG markers of the influence of top-down, context-based prediction on bottom-up acoustic processing also correlated with behavior. These findings help characterize the relationship between brain and behavior by comprehensively linking hierarchical indices of neural speech processing to language comprehension metrics.

**SIGNIFICANCE STATEMENT:** Acoustic and linguistic features of speech have been shown to be consistently tracked by neural activity even in noisy conditions. However, it is unclear how signatures of low- and high-level features covary with one another and relate to behavior across these listening conditions. Here, we find that linguistic (phonetic feature and word probability-based feature) processing is affected by noise more than low-level acoustic feature processing. We also find that behavioral performance is associated with acoustic, phonetic, and lexical surprisal tracking, and that these associations depend on background noise levels. These results extend our understanding of how various speech features are comparatively reflected in electrical brain activity and how they relate to perception in challenging listening conditions.

## INTRODUCTION

Given the importance of speech communication in human life, tremendous amounts of research have focused on characterizing the neurophysiology of language comprehension (Hickok, 2015). This research has revealed a network of brain areas that are functionally specialized for processing different hierarchical levels of speech and language (Hickok & Poeppel, 2007). For example, work has shown that low level acoustic and spectrotemporal features of speech are chiefly processed in early auditory cortex (de Heer et al., 2017), with various phonological features being processed in secondary areas such as the superior temporal gyrus (Hamilton et al., 2018; Mesgarani et al., 2014) and some prefrontal areas (Burton, 2009; de Heer et al., 2017), and meaning being represented across large areas of cortex (Anderson et al., 2017; Huth et al., 2016; Pereira et al., 2018).

While much of the above knowledge has been obtained from functional neuroimaging and invasive recordings in neurosurgical patients, parallel efforts have been made to obtain noninvasive magneto- and electrophysiological (MEG/EEG) markers reflecting the hierarchical processing of speech. This includes modeling how EEG and MEG track the amplitude envelope of natural speech (Lalor & Foxe, 2010) and how neural responses reflect the spectrotemporal (Daube et al., 2019; Di Liberto et al., 2015), phonetic (Di Liberto et al., 2015), phonotactic (Brodbeck, Hong, et al., 2018; Di Liberto et al., 2019; Gwilliams et al., 2022), lexical (Dou et al., 2025; Heilbron et al., 2022), prosodic (Teoh et al., 2019), and semantic (Broderick et al., 2018; Heilbron et al., 2022) features of natural speech.

Some advantages of EEG are that it is significantly cheaper and easier to use in applied research in different cohorts (Peck et al., 2021; Salisbury et al., 2002). Consequently, there has been considerable interest in exploring how different EEG markers of speech processing reflect speech intelligibility (Verschueren et al., 2021) and language comprehension (Ahissar et al., 2001; Broderick et al., 2022). Many of these studies altered the intelligibility or comprehensibility of speech by adding background noise (Iotzov & Parra, 2019; Schmitt et al., 2022; Zhang et al., 2023) or by degrading the speech signal itself (Karunathilake et al., 2023; Viswanathan et al., 2021). For example, some work has shown that cortical tracking of the speech envelope decreases as noise levels increase (Etard & Reichenbach, 2019; Lesenfants et al., 2019; Vanthornhout et al., 2018; Zou et al., 2019), although others have suggested such tracking remains robust until the background noise is more than twice as loud as the speech it masks (Ding & Simon, 2013). Meanwhile, experiments that have explored EEG indices of semantic processing appear to show a strong correlation with speech intelligibility and/or understanding (Broderick et al., 2018). Very few studies, however, have systematically explored how EEG indices of both low- and high-level speech processing covary across different levels of speech comprehension (Mesik et al., 2021; Sohoglu & Davis, 2020; Strauß et al., 2022; Yasmin et al., 2023). This is the goal of the present study.

Our study explores how EEG markers of speech envelope, spectrogram, acoustic onset, phonetic feature, and lexical surprisal vary with subjective measures of the percentage of words heard (PWH, as a proxy for intelligibility) and objective measures of comprehension across different background noise conditions. We hypothesize that EEG measures of higher-level processing (e.g., lexical surprisal and phonetic features) will more strongly correlate with behavior than lower-level measures (e.g., spectrogram). We also test how listeners exploit linguistic context to process noisy speech and how that effect might manifest in EEG. Here, we leverage a previously introduced measure of predictive speech perception that quantifies how the tracking of low-level speech features varies as a function of the context-based semantic content of that speech (Broderick et al., 2019), but this time using lexical surprisal. We hypothesize that this measure strengthens for speech in moderate levels of noise (when speech is still intelligible) relative to speech in quiet, before declining at high levels of background noise (when speech is no longer intelligible). With this study we seek to extend our understanding of the hierarchical processing of continuous speech under a range of realistic listening conditions.

## MATERIALS AND METHODS

### Participants

28 healthy adults (9 males, 18-35 years old [mean = 22.08, SD = 4.63]) participated in this study. One participant was excluded due to an insufficient amount of data and two were excluded due to technical issues, resulting in a dataset of 25 participants. Each participant provided written informed consent and reported having normal hearing, normal or corrected-to-normal vision, English as their first and main language, and no history of neurological disorders. Participants were also compensated for their participation. All procedures were approved by the University of Rochester Research Subjects Review Board.

### Stimuli and experimental procedure

Participants listened to 70 minutes of *A Wrinkle in Time* by Madeleine L’Engle which was read by an American female speaker. The stimulus was presented across 70 trials, each being one minute in duration. Each trial was presented at one of five noise levels: quiet (no noise) and +3 dB, −3 dB, −6 dB, and −9 dB signal-to-noise ratios (SNRs). The background noise was spectrally matched stationary noise, which was estimated from the clean speech using a 46^th^ order forward linear predictive model (Crosse, Di Liberto, & Lalor, 2016; Ding & Simon, 2013). Prediction order was defined based on the audio sampling rate as p_order_ = (Fs/1000)+2, to capture vocal tract and non-vocal tract related contributions to speech signals (Atal & Hanauer, 1971). There were 14 minutes’ worth of audio for each of the five noise conditions. The storyline was preserved from trial to trial, but the conditions were pseudorandomized such that no noise level occurred consecutively. Participants estimated the percentage of words they heard (on a scale of 0-100%) and answered two multiple choice comprehension questions after each trial. The comprehension questions used here were the same questions used from a previous study (Maddox & Lee, 2018), except we presented only two out of the four original questions created for each trial. The stimuli were presented through Sennheiser HD650 headphones at a sampling rate of 44.1 kHz using Psychtoolbox (Kleiner et al., 2007) and custom MATLAB scripts (MathWorks, 2019).

### Data acquisition and preprocessing

EEG data were recorded from 128 scalp electrodes (plus two mastoid channels that were not analyzed in this work). The data were acquired at a 1024 Hz sampling rate with the BioSemi Active Two system. The data were preprocessed using the PREP pipeline and its default parameters (Bigdely-Shamlo et al., 2015). This pipeline first detrended the data at 1 Hz, then removed the 60 Hz line noise. Afterwards, robust re-referencing was applied which allows the data to be referenced to an average of all channels except those contaminated with noise. This function identifies and interpolates noisy channels in an iterative manner such that the re-referencing itself is not affected by the noise. The cleaned data was then low pass filtered at 8 Hz, using a Chebyshev Type II filter with an 8.5 Hz cutoff frequency and an 80 dB stopband attenuation. Next, the data were epoched and independent component analysis (ICA) was applied using EEGLAB’s *picard* function (Delorme & Makeig, 2004; Pion-Tonachini et al., 2019) to remove muscle and eye artifacts. Lastly, the data were downsampled to 128 Hz.

### Speech stimulus characterization

Speech is organized in a hierarchical manner where sounds can form syllables, syllables form words, words form sentences, and so on. To assess how our brains might concurrently process speech across levels of this hierarchy, we chose to model EEG responses to speech based on several different representations of that speech, all of which were computed using the clean (no noise) versions of each trial.

#### Envelope

We first calculated the speech envelope, a well-established feature shown to be robustly tracked by cortical activity (Aiken & Picton, 2008; Destoky et al., 2019; Di Liberto et al., 2015; Ding & Simon, 2013; Etard & Reichenbach, 2019; Lalor & Foxe, 2010; Nourski et al., 2009; Pasley et al., 2012) and to be important for speech recognition and intelligibility (Ahissar et al., 2001; Drullman et al., 1994; Shannon et al., 1995). The speech signal was first lowpass filtered at 20 kHz (22.05 kHz cutoff frequency, 1 dB passband attenuation, 60 dB stopband attenuation). The broadband speech envelope was calculated using a gammachirp auditory filterbank to mimic the filtering properties of the cochlea (Irino & Patterson, 2006). This filterbank was used to filter the speech into 16 bands from 250 Hz to 8 kHz with an equal loudness contour (essentially creating a spectrogram). Lastly, we calculated the absolute value of the Hilbert transform for each frequency band and averaged this result across all 16 bands.

#### Acoustic onsets and spectrogram

We chose to model two additional acoustic features, acoustic onsets and spectrogram, which were shown to be reflected in cortical activity above and beyond the speech envelope (Brodbeck et al., 2020; Di Liberto et al., 2015; Sohoglu & Davis, 2020). Acoustic onsets were approximated by computing the first derivative of the speech envelope and then half-wave rectifying the result (Hertrich et al., 2012). A 16-band spectrogram was computed using the same filterbank and parameters as the speech envelope, just without the final averaging step.

#### Phonetic features

To calculate phonetic features, the Montreal Forced Aligner (McAuliffe et al., 2017) was first used to partition and time align each word in the story into phonemes according to the International Phonetic Alphabet for American English. Then, each phoneme was linearly mapped onto a set of 19 binary phonetic features based on the University of Iowa’s phonetics project (http://www.uiowa.edu/~acadtech/phonetics/english/english.html/).

#### Lexical surprisal

Lastly, we calculated the surprisal of each word based on its preceding context using the Transformer-XL model (Dai et al., 2019). This model contains a recurrence mechanism that allows it to build and reuse memory from previous segments and learn longer-term dependencies, while preserving the temporal information of previous word embeddings. This model was chosen because it can predict the probability of an upcoming word using the context from all preceding words. The softmax of the values from the output layer of the model were taken to estimate the probability of each word, and the negative log of a word’s probability was computed to estimate its lexical surprisal (Dai et al., 2019).

### Modeling the relationship between speech features and EEG responses

One goal of the present study was to determine how neural representations of individual speech features change with SNR and how those changes relate to comprehension and what participants reported hearing. To index the fidelity of individual speech feature representations, we used a forward encoding model (in the form of a linear filter or kernel, sometimes known as a temporal response function) which describes the transformation from the speech features to the EEG responses recorded at each electrode. Here, acoustic onsets, the spectrogram, phonetic features, and lexical surprisal were modeled. An additional vector with impulses placed at each word’s onset was included to capture any acoustic related onset responses at word boundaries that the acoustic onset predictor may have missed. The forward encoding models can be expressed as follows:

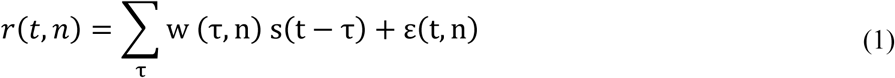

where s(t − τ) is the speech feature of interest, 𝑟𝑟(𝑡𝑡, 𝑛𝑛) is the neural response on each channel, *n*, w (τ, n) is the temporal response function to be estimated, and ε(t, n) is the residual recorded EEG not explained by the model, which is assumed to be Gaussian.

Because our five different speech features of interest (spectrogram, acoustic onsets, phonetic features, lexical surprisal, and word onsets)—like almost all speech features—are correlated with each other in time, we wanted to use an approach where we could identify the unique contribution that each speech feature made to the resulting EEG responses. To do this, we used a partial correlation approach. Specifically, our strategy involved first modeling the data from each of our experimental (SNR) conditions *separately* using each of our five speech features. This resulted in five TRF models for each condition, calculated from training data. We then used the five speech features from left-out trials, and the corresponding TRF models, to generate five separate predictions of the EEG responses for those trials.

Then, to calculate how much each speech feature uniquely contributed to the recorded brain activity, we computed a partial correlation between the real EEG and the predicted EEG from each speech feature, while accounting for the predicted EEG from all other speech features. For example, to assess how much the acoustic onsets [ons_A_] uniquely contributed to the EEG responses, we concatenated the predicted EEG from the other four feature models (spectrogram [sgram], phonetic features [fea], surprisal [surp], word onsets [ons_W_]) into a matrix, Z. We then used MATLAB’s partialcorr (X,Y,Z) function (Fisher, 1924) to determine the partial (Pearson’s) correlation between the actual EEG (X) and the predicted EEG from the model trained on the acoustic onsets (Y), while controlling for the activity that can be explained by all other features (Z). We repeated this process to identify the unique contribution of each speech feature, while controlling for all other speech features.

In terms of training and testing the models, nested leave-one-out cross-validation and ridge regression were used. Each speech feature was normalized between 0 and 1 by dividing by its maximum value, and the EEG data were z-scored. The features and EEG were partitioned into train and test sets. The stimuli were lagged from −100-700ms to capture both short and long latency responses to acoustic and linguistic features. Given the amount of data, models were validated on the clean condition and tested on the remaining noise conditions. Specifically, cross-validation was conducted on the *clean* condition to select the optimal regularization (ridge) parameter, *λ*, which ranged from 10^-1^–10^8^. We identified the regularization parameter that resulted in the highest prediction accuracy for each individual test fold. We then selected the parameter that produced the highest prediction accuracy most often (across all test folds) so that we could use one parameter to train the models for each condition. Using the same parameter for all folds and conditions (within a participant) allowed for a fairer comparison of model performance since it results in the predictions across trials being on the same scale for each participant. This method was particularly important for the low SNR conditions, as the models would be more prone to overfitting, especially when training on multidimensional feature spaces.

Our primary analysis aimed to assess how different hierarchical speech features related to EEG activity across listening conditions. However, given its prevalence in the literature and relative ease of modeling, we also examined how speech envelope tracking varied with SNR. To do this, we used backward modeling to reconstruct an estimate of the speech envelope from the recorded EEG for each trial. The backward model or decoder, g(τ, n), described the transformation from lagged EEG responses at all electrodes, r(t + τ, n), to an estimate of the speech envelope, 𝑠𝑠^(𝑡𝑡). As detailed elsewhere (Crosse, Di Liberto, Bednar, et al., 2016; Crosse et al., 2021), the modeling procedure can be expressed as:

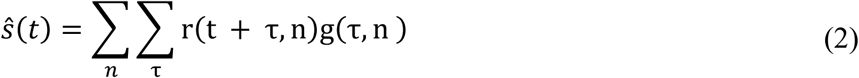

Similar to the forward models, this analysis was conducted separately for each experimental (SNR) condition using 14-fold leave-one-out cross-validation. First, the stimuli and responses were normalized and then partitioned into train and test sets. Here the EEG data were lagged, τ, from −100–300ms to focus on short latency responses to the speech envelope, s(t). Cross-validation and regularization parameter (10^-1^–10^8^) selection followed the same procedure as described above for forward modeling. A decoder was trained using the training set and selected regularization parameter to reconstruct an estimate of the speech envelope. After testing the decoder on held-out data, model performance was assessed by computing the Pearson correlation between the actual speech envelope and the reconstructed speech envelope. This model performance, also known as reconstruction accuracy, was averaged across folds and participants (Figure 1).

**Figure 1.**
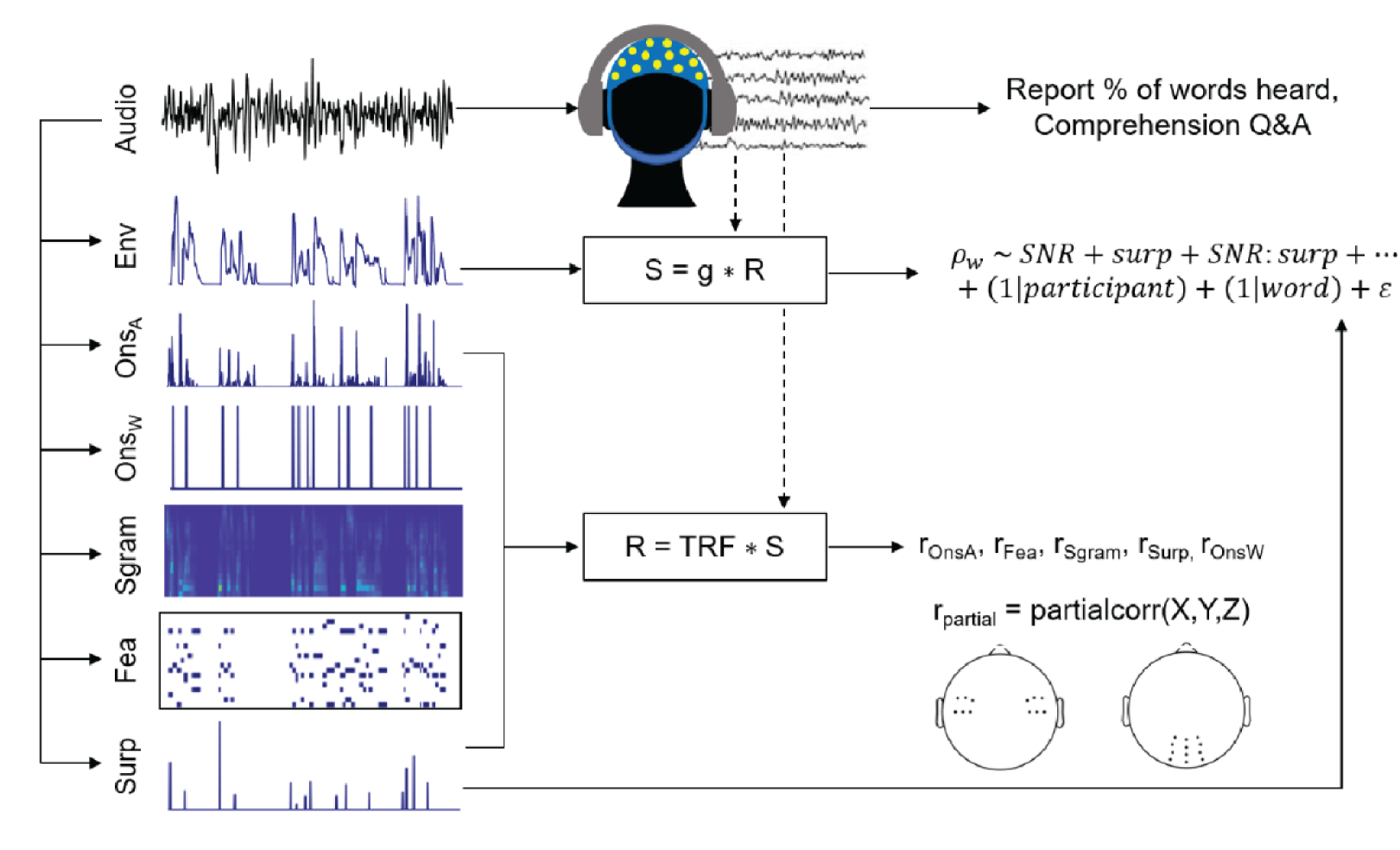
Methods. EEG data were recorded while participants listened to an audiobook in different levels of noise. They also reported the percentage of words they heard (%) and answered comprehension questions (Q&A). Forward modeling (with a temporal response function, TRF) was used to estimate EEG responses (R) from the clean representations of the speech features (S), which were acoustic onsets (onsA), word onsets (onsW), spectrogram (sgram), phonetic features (fea), and word surprisal (surp). Model performance (r) was assessed by calculating the partial correlation between the actual EEG and predicted EEG from one feature’s model, while controlling for the predictions generated by all other features. These partial correlations were averaged across the selected electrode channels marked in the head plots. Backward modeling (with a decoder, g) was also used to reconstruct an estimate of the clean speech envelope (env, S). Model performance (reconstruction accuracy, *ρw*) was assessed by calculating the correlation between the original speech envelope and the reconstructed speech envelope. A linear mixed-effects model (LME) was then used to determine the influence of word surprisal on envelope reconstruction accuracy.

### Assessing the role of lexical context on acoustic encoding in different levels of background noise

Another major goal of the present study was to test how listeners might rely more on lexical context to—perhaps predictively—encode speech that is masked by moderate levels of background noise. To do this, we used a variant of a recently introduced approach that involved quantifying how the tracking of low-level speech features varies as a function of the context-based semantic content of that speech (Broderick et al., 2019). In our variation, we used a linear mixed-effects (LME) model to explore the extent to which word predictability in the form of lexical surprisal as opposed to semantic similarity influenced how the envelope of that word was reflected in EEG and how that influence changed across SNRs. By using an LME model in this way, one can measure the relationship between the main variable(s) of interest while controlling for variability caused by random factors. We used the *lmerTest* package in R (Kuznetsova et al., 2017) to model the following equation:

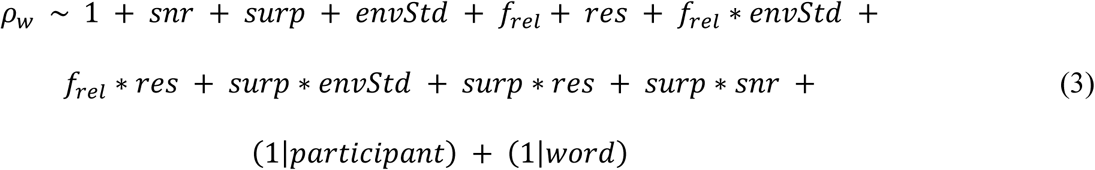

The dependent variable is word reconstruction accuracy which was calculated as the Spearman’s correlation between the actual word envelope and the predicted word envelope for the first 100 ms of each word. As detailed below, the independent variables are SNR, lexical surprisal (*surp*), envelope variability (*envStd*), relative pitch (*f_rel_*), resolvability (*res*), and various interactions such as the interaction between surprisal and SNR.

Envelope variability, relative pitch, and resolvability were selected as nuisance regressors. This was because these measures can correlate with surprisal, with one another, and with envelope tracking; so, following Broderick and colleagues, they are included here and as interactions to ensure that they aren’t inherently contaminating the lexical surprisal effects (Broderick et al., 2019). Envelope variability is represented as the standard deviation of the speech envelope. Relative pitch is pitch normalized to the vocal range of the speaker (Tang et al., 2017) and resolvability measures whether a sound’s harmonics are processed between distinct (resolved) or within the same (unresolved) filters of the cochlea (Shackleton & Carlyon, 1994). Prosodic cues such as relative pitch and resolvability can uniquely predict EEG activity even after accounting for other acoustic and phonetic features (Teoh et al., 2019) which was yet another reason to include them in this LME model. Relative pitch was extracted using Praat (Boersma & Weenink, 2013). Once the software estimated absolute pitch, this result was then z-scored, resulting in relative pitch. Resolvability was extracted using custom scripts based on a model of the human auditory periphery (McDermott & Simoncelli, 2011; Teoh et al., 2019).

The LME model also included by-word and by-participant random intercepts as some words may be easier to reconstruct than others and some participants may, on average, have higher reconstruction accuracies than other participants (based on cortical folding and other biophysical factors). No random slopes were included, as they caused the model to not converge (calculate a solution) even with the addition of an optimizer (additional functions the model can use to reach a solution (Brown, 2021)). Like Broderick and colleagues, the speech envelope and the nuisance regressors were calculated for the duration of the trials. Afterward, word reconstruction accuracy was calculated over the first 100 ms following each word’s onset and the nuisance regressors were segmented over the same interval (Broderick et al., 2019). There were 11,748 words total with an average length of 292.3 ms and a standard deviation of 182.3 ms.

### Statistical analysis

All statistical analyses were performed in R (version 4.5.0) and in MATLAB R2024b (MathWorks, 2024). Due to the skewed distribution of the behavioral scores, comparisons were calculated using a nonparametric Friedman test, followed by a Wilcoxon Rank Sum test with Holm correction. Pairwise tests were also used to determine differences in envelope reconstruction accuracy and EEG partial correlations between conditions. Permutation testing was performed to test if the partial correlation coefficients were greater than chance. The predicted EEG trials were shuffled, and the partial correlation was computed between the actual EEG (in its correct order) and the predicted EEG (which was out of order). This was repeated 1,000 times for each participant. The null partial correlations were averaged across folds, electrodes (12 parietal electrodes for the surprisal model and 12 frontotemporal electrodes for all other feature models, all selected *a priori*, Figure 1), and participants, and we assessed the proportion of permuted coefficients that were greater than the actual group-averaged coefficients. Lexical surprisal is known to be reflected on more parietal areas of the scalp and displays more N400-like responses, and thus the surprisal partial correlations were averaged across the selected parietal electrodes (Broderick et al., 2022). All other features were averaged across the frontotemporal electrodes as those regions have been shown to reflect cortical auditory responses (Di Liberto et al., 2015).

Permutation testing was also performed to find if the speech envelope reconstruction accuracies were greater than chance. A null model was created for each SNR by shuffling the speech envelope between trials and calculating a new model with each shuffle. This procedure, including regularization, was repeated 1,000 times. The null reconstruction accuracies were averaged across folds and participants and then we assessed the proportion of reconstruction accuracies that were greater than the actual reconstruction accuracies.

One question of interest was how the neural representation of each speech feature varied with noise. To test this, the partial correlations for each trial were used in an LME model. This model (Equation 4) included SNR, speech feature type, and the interaction between the two as fixed effects, and a by-participant random intercept.

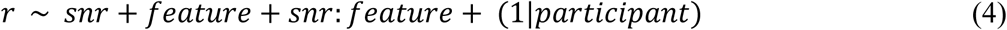

To compare the changes in partial correlations across SNRs between each feature’s model (essentially comparing the slopes of the SNR by feature interaction), we computed their estimated marginal means and performed pairwise comparisons with Tukey Honestly Significant Difference (HSD) correction using the *emmeans* package in R (Lenth et al., 2019).

Generalized linear mixed models (GLMM) were also used to model the relationships between behavior and partial correlations/reconstruction accuracies (Equation 5) and the surprisal*SNR coefficients from Equation 3 (Equation 6).

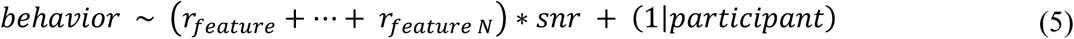

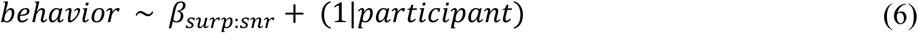

Equations 5 and 6 utilized ordered beta distributions with a logistic link function (which can support the skewed and bounded nature of our behavioral data (Smithson & Verkuilen, 2006)) and the template model builder (glmmTMB) function. The *glmmTMB* package allowed us to model the ordered beta distribution with the addition of random effects (Brooks et al., 2017).

Each mixed-effects model in this study was calculated using z-scored predictors, the default parameters which included fitting the models with restricted maximum likelihood (REML), and Satterthwaite’s method for the t-tests. Marginal and conditional R^2^ values are provided via the *performance* package (Lüdecke et al., 2021) to indicate the variance explained by fixed effects alone (R^2^_m_) and by both fixed and random effects (R^2^_c_) (Nakagawa et al., 2017). The remaining packages used to perform statistical analysis and plotting in R were completed using *ggeffects* (Lüdecke, 2018), *ggplot2* (Wickham, 2011), and *patchwork* (Pedersen, 2019).

## RESULTS

### Speech perception decreases as SNR decreases

Behavioral scores were collected to show how participants’ perception of the story changed as listening conditions became more challenging. After each one-minute-long trial, participants estimated the PWH on a scale from 0-100% and answered two multiple-choice comprehension questions, each of which had four possible answers. As expected, there was a significant reduction in the PWH (*Χ*^2^(4) = 98.894, p < 2.20 x 10^-16^) and comprehension scores (*Χ*^2^(4) = 83.44, p < 2.20 x 10^-16^, Friedman test followed by Wilcoxon rank sum test with Holm correction, Figure 2) with the addition of noise. After the experiment, participants reported hearing very few words in the −9 dB SNR condition but attempted to use context clues from previous trials to answer the questions in the −9 dB SNR trials. To control for this potential confound in our measure of comprehension, we used a procedure similar to Orf and colleagues, where nine additional volunteers answered the comprehension questions for each trial without listening to the story (Orf et al., 2022). Based on the performance of those new participants, we determined a new empirical chance level of 28% rather than 25%. All participants performed above chance in the quiet, +3 dB SNR, and −3 dB SNR conditions. Comprehension scores were not above chance for five participants in the −6 dB SNR condition (p ≥ 0.066) and 13 participants in the −9 dB SNR condition (p ≥ 0.066). Although participants reported hearing fewer words in the +3 dB SNR condition compared to speech in quiet, they performed similarly in terms of comprehension (p = 1.000, S1 Table).

**Figure 2.**
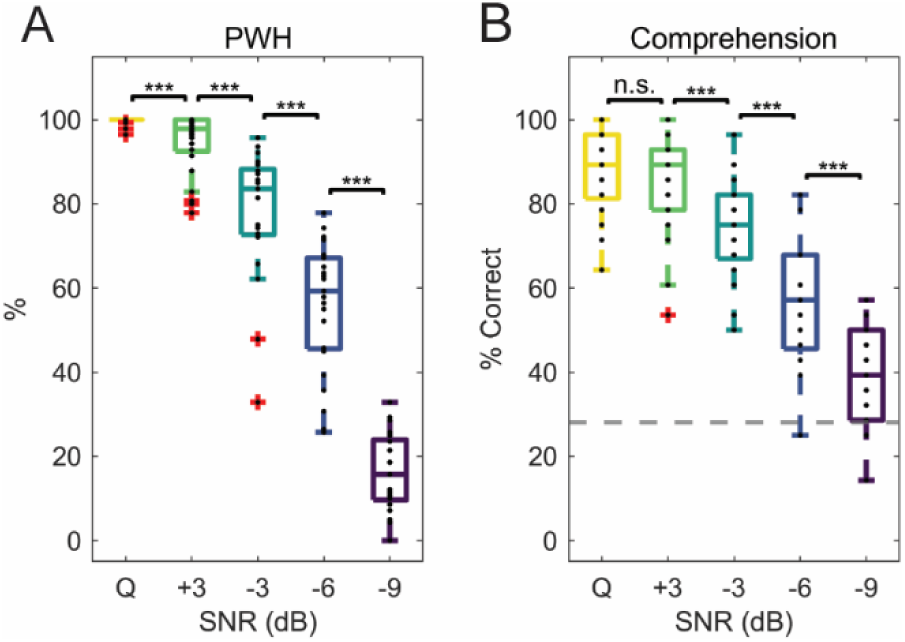
Behavioral results. (A) Average percentage of words participants reported hearing in each condition. (B) Average percentage of correctly answered comprehension questions for each condition. The dashed gray line is the chance level at 28% which was calculated using a separate set of participants who answered the comprehension questions without listening to the audiobook. For both plots, significance is indicated by * if p < 0.05, ** if p < 0.01, and *** if p < 0.001 using pairwise Wilcoxon rank sum tests. The black dots in each plot correspond to individual participants and the box and whiskers depict the median across participants, in addition to the 75^th^ and 25^th^ percentiles.

### Hierarchical speech feature encoding declines across SNRs

Many EEG/MEG speech studies have focused on modeling brain responses to isolated speech features, perhaps most frequently the speech envelope. Recent work, however, is increasingly using various methods to model acoustic and linguistic features simultaneously (Brodbeck, Hong, et al., 2018; Brodbeck, Presacco, et al., 2018; de Heer et al., 2017; Gillis et al., 2021; Heilbron et al., 2022; Verschueren et al., 2022) to find how the brain uniquely encodes a feature of interest while accounting for others, as some features may be correlated and explain similar neural activity (Daube et al., 2019). Since few studies have modeled the encoding of simultaneous features in challenging listening conditions (Brodbeck et al., 2020; Karunathilake et al., 2023; Sohoglu & Davis, 2020), we were interested in how a range of acoustic and linguistic features were encoded in noise.

To test this, we quantified the unique contribution that each speech feature makes to the EEG responses using partial correlation calculations. Specifically, separate models were fit on the acoustic onset, speech spectrogram, phonetic feature, word surprisal, and word onset features. Those models were tested on left out data to predict unseen EEG. The partial correlation was then calculated between the actual EEG and the predicted EEG for one feature while controlling for the effects of all others. The acoustic onset, spectrogram, phonetic feature, and word onset model partial correlations were averaged across 12 frontotemporal electrodes (Di Liberto et al., 2015) and the surprisal partial correlations were averaged across 8 parietal electrodes (Broderick et al., 2022) plus 4 additional electrodes in the same area so that both analyses consisted of 12 electrodes (electrodes shown in Figure 1). Model partial correlation coefficients were above chance for each SNR (p ≤ 0.021, permutation tests) except for the phonetic feature model in the −9 dB SNR (p = 0.157) and the word onset model in the −6 dB and −9 dB SNRs (p = 0.146, p = 0.333). Unlike the other features, there were no significant differences in the average partial correlation coefficients for the spectrogram model between SNRs (Figure 3A, within feature paired tests with Tukey HSD correction).

**Figure 3.**
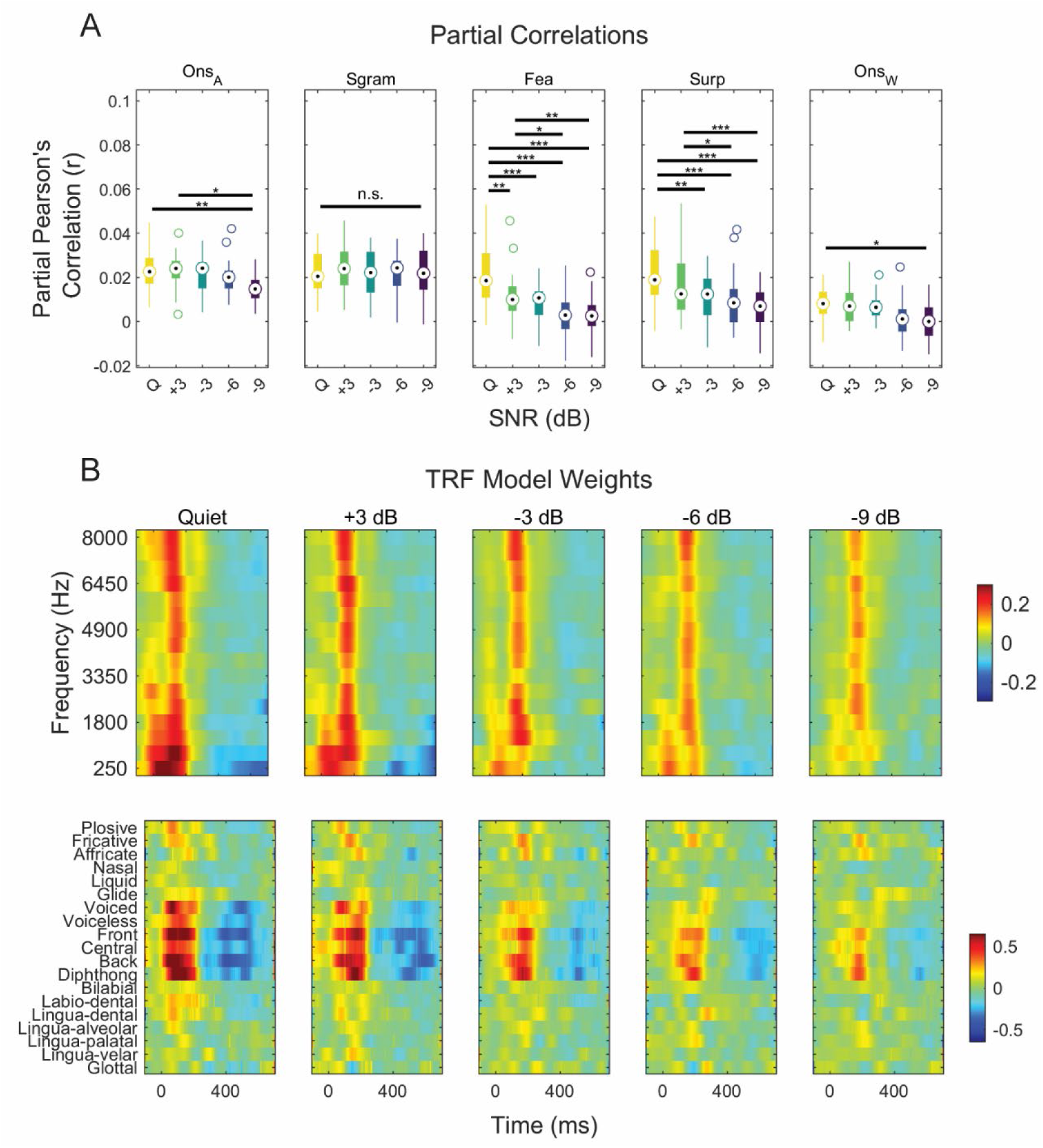
Forward modeling results. (A) Partial correlations were calculated based on actual EEG and predicted EEG from models based on acoustic onsets, spectrogram, phonetic features, surprisal, and word onset using MATLAB’s partialcorr function. Significance is indicated by * if p < 0.05, ** if p < 0.01, and *** if p < 0.001 using within-feature pairwise t-tests with Tukey’s HSD correction. (B top) Spectrogram and (B bottom) phonetic feature TRF model weights across the experimental conditions. The model weights were averaged across the 12 frontotemporal electrodes depicted in Figure 1.

A subsequent LME model analysis (Equation 4) was conducted to examine how the neural representation of different speech features decreases with increasing levels of noise. This helps account for the fact that EEG partial correlation coefficients can vary greatly between participants (based on, for example, cortical folding or skull/scalp geometry). Since we were interested in trends across successive noise levels, SNR was represented using a linear polynomial contrast which tests for a monotonic trend across these ordered noise levels. In this case, the order was set from quiet to −9 dB so that the results would be easily comparable to the partial correlation results shown in Figure 3A. Due to the large number of variables included in the model, a reference level was also needed. The acoustic onset category of the *feature* variable was set as the reference level using a dummy coding scheme. Since the reference level also serves as the model intercept when dummy coding, every other category of the *feature* variable represents the adjustment to that intercept. Similarly, the interaction coefficients can be interpreted as an adjustment to the *SNR* variable because, due to the dummy coding scheme, it represents how partial correlations for the acoustic onset model varied across SNRs (Brown, 2021).

The LME model (R^2^_m_ = 0.086, R^2^_c_ = 0.111) results are shown in Table 1. The raw fixed effect outputs are listed in Section 1. Section 2 reports the estimated marginal slopes for each feature by SNR interaction, along with statistics testing whether each slope differs from zero. Finally, Section 3 reports Tukey HSD corrected pairwise comparisons between the interaction effects from Section 2. With acoustic onsets as the reference level and SNR coded using a linear polynomial contrast, the main effect coefficients quantify overall differences between acoustic onsets and every other feature evaluated at the mean SNR. As such, each feature’s main effects differed significantly from the acoustic onset reference, indicating feature-specific differences in partial correlation strength at the mean SNR (p = 0.010 for spectrogram and p < 2.22 x 10^-16^ for phonetic features, surprisal, and word onset).

**Table 1.**
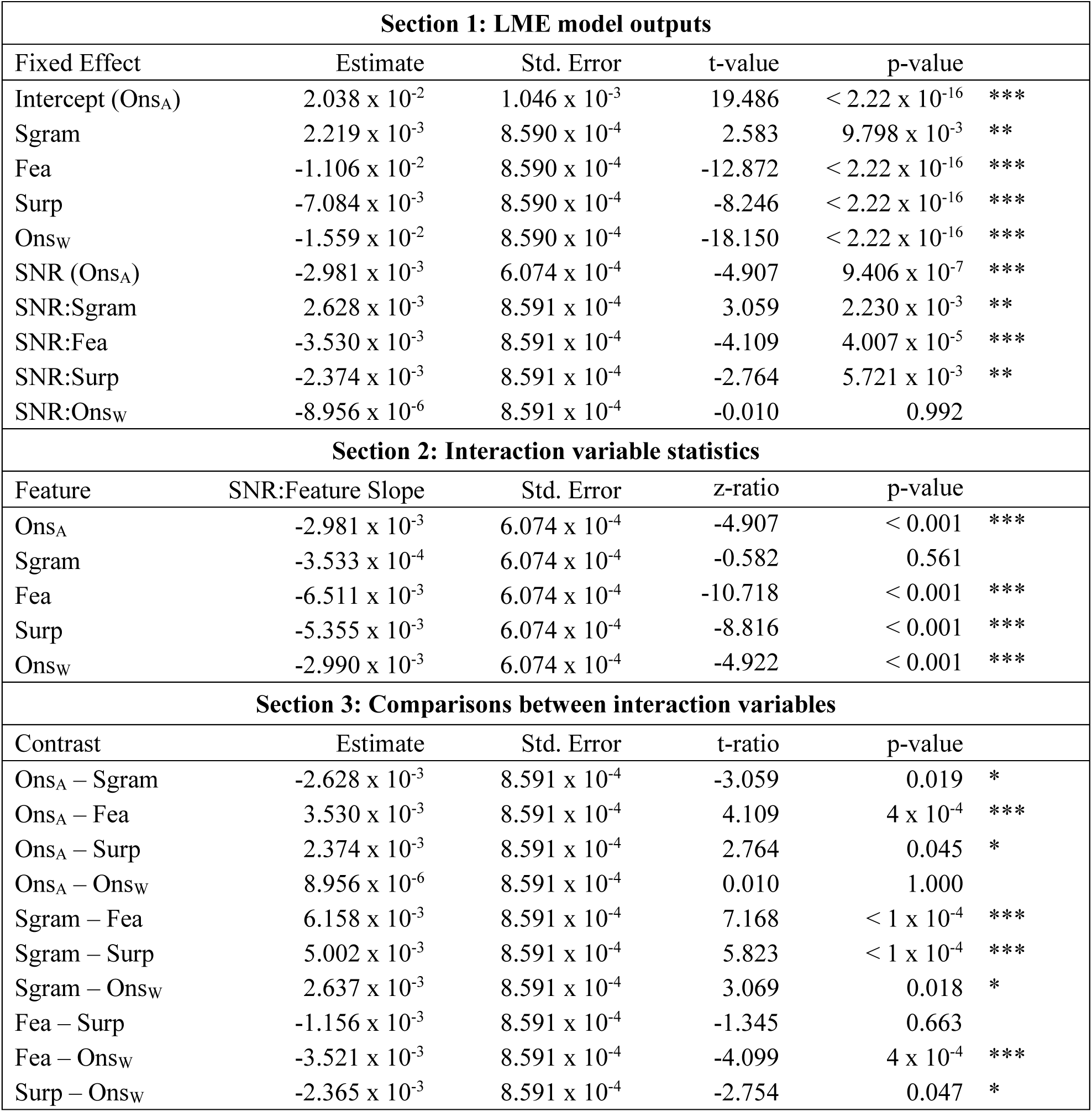
LME model results which show how the partial correlation coefficients changed across SNRs in addition to post-hoc contrasts between each feature.

We were particularly interested in the interaction effects, since we wanted to know how the partial correlations for each model varied across SNRs. EEG predictions based on the speech spectrogram had the flattest slope across noise conditions (β = −3.533 x 10^-4^, p = 0.561, Table 1, Section 2) suggesting that after controlling for all other features, the speech spectrogram exhibited the least change in how it was reflected in the EEG as noise levels increased. Another notable finding was that the changes in partial correlation coefficients for both the phonetic feature (β = − 6.511 x 10^-3^) and surprisal (β = −5.355 x 10^-3^, Table 1, Section 2) models were similar to one another (p = 0.663), yet different from all other models (p < 0.05, Table 1, Section 3). The partial correlation coefficient trends for the phonetic feature and surprisal models decreased at a faster rate compared to the other features which suggests that phonetic feature and surprisal representations were more sensitive to noise. Altogether, the LME model analysis and post-hoc comparisons showed that neural representations of the speech spectrogram were least affected by noise and that the higher-level speech feature representations were more sensitive to increasing levels of noise than the lower-level acoustic feature representations.

Another way to examine the effect of SNR on speech feature encoding is to visualize the TRF models themselves. In fact, recent work has shown that the amplitude and latency of acoustic onset and semantic TRFs are influenced by speech SNR (Yasmin et al., 2023). Given that the encoding of specific frequency bands or phonetic properties may decrease differentially with noise, we examined the change in the spectrogram and phonetic feature TRF weights at each SNR. Since the main positive peaks are seen within the first 200 ms, the model weights were averaged from 0-200ms to better examine how the TRF weights changed across that time range (S1 Figure).

In quiet, we saw the strongest TRF weights in the lowest frequency bands, suggesting the importance of low frequency spectral tracking when no noise is present. Notably, model weights from 250-1,800 Hz seem to diminish in the −6 dB and −9 dB SNR conditions compared to quiet (p < 0.05 and p < 0.001, pairwise tests with Tukey correction). Those same model weights were different in −9 dB compared to the +3 dB condition (p < 0.01, paired tests with Tukey correction, Figure 3B top, S1 Figure). These results support the importance of lower frequencies in speech encoding which is reasonable given evidence that this range is useful under marginally intelligible conditions (Bröhl & Kayser, 2021) and when supplementing listening at higher noise levels (Chang et al., 2006; Turner et al., 2004). However, it is important to note that although there was a reduction in spectrogram TRF weights across SNRs, the spectrogram partial correlation coefficients did not show a similar pattern. This is likely due to the TRF being fit on the individual feature, whereas the partial correlation analysis controlled for the contribution of all other features. Therefore, the TRF weight results should be interpreted with caution.

In the phonetic feature case, TRF amplitudes decreased for the majority of features as noise levels increased (Figure 3B bottom). Voicing and vowel backness features (central, back, and diphthong) on average remained largely intact in the +3 dB condition compared to quiet (p > 0.05 for each, paired t-test with Tukey correction, S1 Figure) but decreased in all other conditions (including the front feature, p < 0.05 for each, paired t-test with Tukey correction, S1 Figure 1, Figure 3B bottom). In the −6 dB SNR condition, only some fricative and vowel backness weights remained whereas all other features here and in the −9 dB SNR condition diminished (Figure 3B bottom). Although TRF weights for some individual features decreased in the +3 dB SNR, we witnessed a larger reduction in TRF amplitudes in the −6 dB and −9 dB SNRs similar to Swaminathan and Heinz who found their greatest lapse in phonetic feature reception around their −5 dB and −10 dB SNRs (Swaminathan & Heinz, 2012).

### Feature-specific brain–behavior relationships change with SNR

One of the primary hypotheses of this study was that EEG signatures of linguistic processing would be more closely related to behavior than EEG measures of low-level acoustic processing, particularly as noise levels increased. To test this, we modeled the relationship between behavioral scores and each speech feature’s partial correlations across SNRs using GLMMs (Equation 5) with logistic link functions. There is one GLMM for each behavioral metric and all partial correlations are included in each model to control for the effects of the other speech features. In this framework, the main effect of each feature reflects the association between partial correlations and behavior averaged across noise levels, whereas the feature by SNR interaction tests whether the strength of this association depends on SNR. SNR was again represented using a linear polynomial contrast to test for a monotonic trend across noise levels. Here, the order of this variable was set from −9 dB to quiet to represent an increase in SNR. Both behavioral metrics were scaled between 0 and 1 and the predictors were z-scored prior to modeling.

S2 Table displays information about each GLMM such as the intercept, fixed effect estimates, standard error for the fixed effect, z-value, and p-value. Figure 4 depicts the main effect of each feature, with significant interactions shown in inset plots. The predicted lines in each subplot were generated from the GLMMs using *ggpredict* (from the *ggeffects* package) and represent the expected behavioral scores for each combination of predictors, averaged over participant-level random effects. Because logistic link functions were used, the fixed effect estimates represent the amount that the log odds of the behavioral scores change given a unit increase in the partial correlation and partial correlation by SNR coefficients.

**Figure 4.**
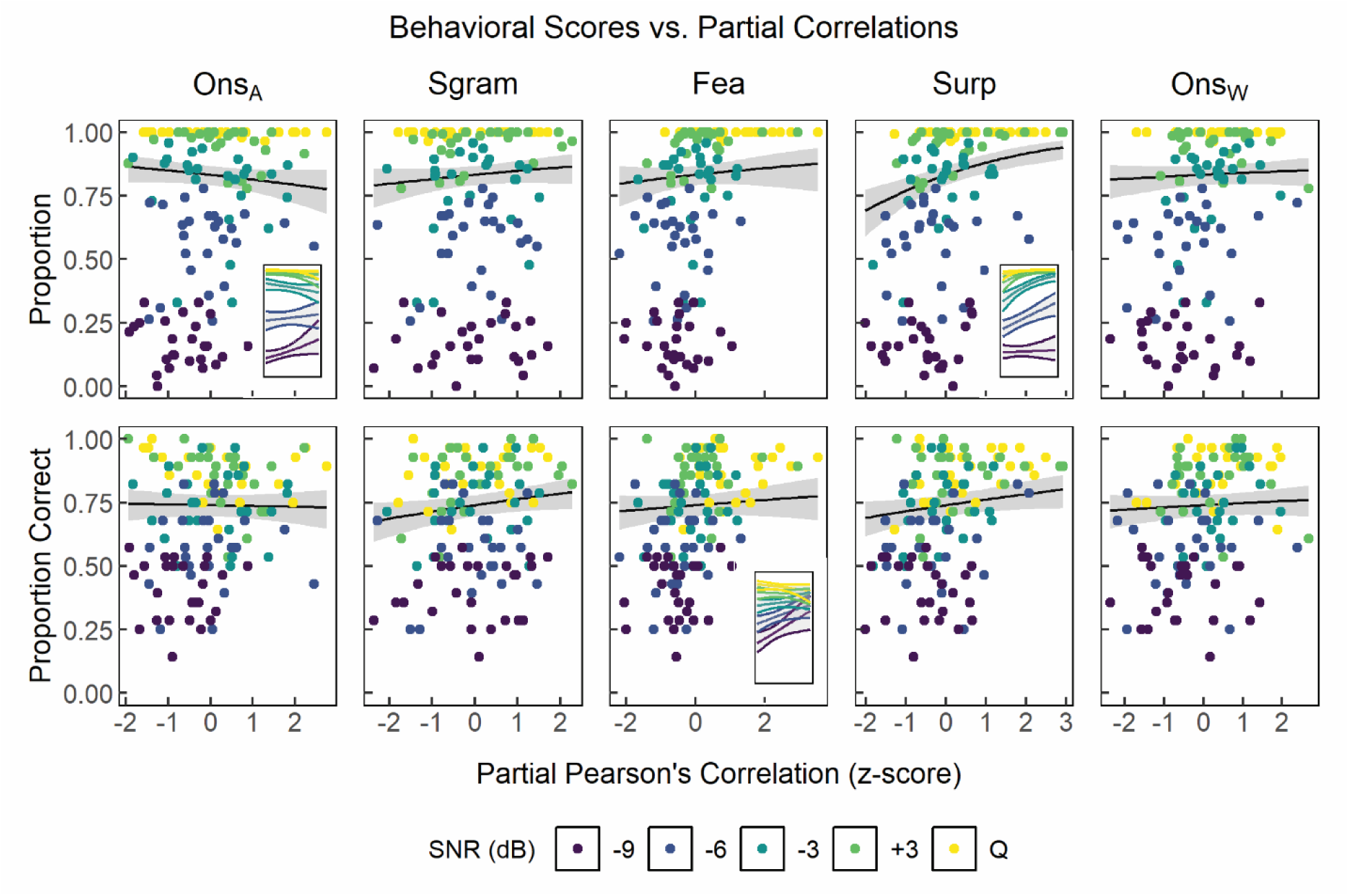
The relationship between the partial correlation coefficients and behavior. GLMMs were used to relate the neural tracking of each speech feature to participants’ PWH (top) and comprehension scores (bottom). Both models were fit using an ordered beta distribution and a logistic link function. The black lines represent GLMM-predicted behavioral scores (via *ggpredict*) that show the expected effect of each speech feature’s tracking while controlling for SNR and tracking of all other features. The shaded grey regions are 95% confidence intervals, and the colored dots represent each participant’s average partial correlation coefficient in each SNR. The inset plots show predicted behavioral scores for features whose relationship with behavior significantly varied across SNRs. Each SNR is represented by a fitted line, with lightly shaded areas indicating the 95% confidence intervals.

The PWH GLMM (R^2^_m_ = 0.944, R^2^_c_ = 0.997) showed robust overall effects of SNR, such that PWH scores increased as SNR increased (β = 2.132, p < 2.22 x 10^-16^). Among the neural tracking predictors, a significant main effect was observed only for surprisal-related neural tracking, indicating that increased neural tracking of lexical surprisal was associated with higher PWH scores (β = 0.398, p = 3.652 x 10^-5^, Figure 4, top). Critically, however, this relationship between behavior and neural tracking varied across noise levels as reflected by a significant surprisal by SNR interaction, demonstrating that the relationship between lexical surprisal tracking and PWH strengthened as SNR increased (β = 0.265, p = 7.067 x 10^-3^). A significant interaction was also observed for acoustic onsets, such that the relationship between acoustic onset tracking and PWH weakened as SNR increased (β = −0.258, p = 4.820 x 10^-3^). Intuitively, this pattern suggests that as listening conditions improve, word report scores become less linked to low-level acoustic onset tracking and increasingly linked to high-level lexical surprisal tracking.

For the comprehension score GLMM (R^2^_m_ = 0.771, R^2^ = 0.986), behavioral performance again increased as SNR increased (β = 0.830, p <0.001). While positive effects of spectrogram (β = 0.128, p = 0.042) and surprisal (β = 0.122, p = 0.046) tracking indicated overall associations with comprehension scores across SNRs, only phonetic-feature tracking exhibited a significant interaction with SNR (β = −0.124, p = 0.033). Phonetic-feature tracking appears to play a more prominent role in supporting comprehension when speech is acoustically degraded, but it becomes less informative as SNR improves. Overall, these results suggest a noise-dependent division of labor across neural feature representations, with PWH score reliance shifting from acoustic to lexical tracking as listening conditions improve, and comprehension score reliance on phonetic-level tracking decreasing as listening conditions improve.

### Envelope reconstruction accuracy maps well to measures of behavior

Numerous studies have shown that cortical activity tracks the temporal modulations of the speech envelope (Ahissar et al., 2001; Aiken & Picton, 2008; Destoky et al., 2019; Di Liberto et al., 2015; Ding & Simon, 2013; Etard & Reichenbach, 2019; Lalor & Foxe, 2010; Nourski et al., 2009; Pasley et al., 2012). Speech envelope reconstruction is a powerful method of indexing speech tracking, given its advantage of using all scalp data to provide a better signal to noise ratio. Yet, it is unknown exactly what information the speech envelope carries because it has been shown to relate to syllabic boundaries (Hertrich et al., 2012; Oganian & Chang, 2019), phonetic feature information (Rosen, 1992), and prosodic cues (Myers et al., 2019). Nevertheless, due to its common usage in speech research and its improved SNR, we were interested in how speech envelope tracking would change across our experimental conditions and how it would relate to behavior apart from the other speech features.

The first step in this analysis was to use a backward modeling approach to determine how speech envelope tracking was affected by different levels of background noise. Speech in quiet and in the +3 dB SNR condition were reconstructed to a similar degree (p = 0.305, paired t-test with Holm correction) and were reconstructed more reliably than all other SNRs (p < 0.05). The − 3 dB SNR and −6 dB SNR accuracies were not significantly different from each other (p = 0.305) but were both different from the −9 dB SNR condition (p = 1.748 x 10^-7^ and p = 2.347 x 10^-6^, respectively, Figure 5A).

**Figure 5.**
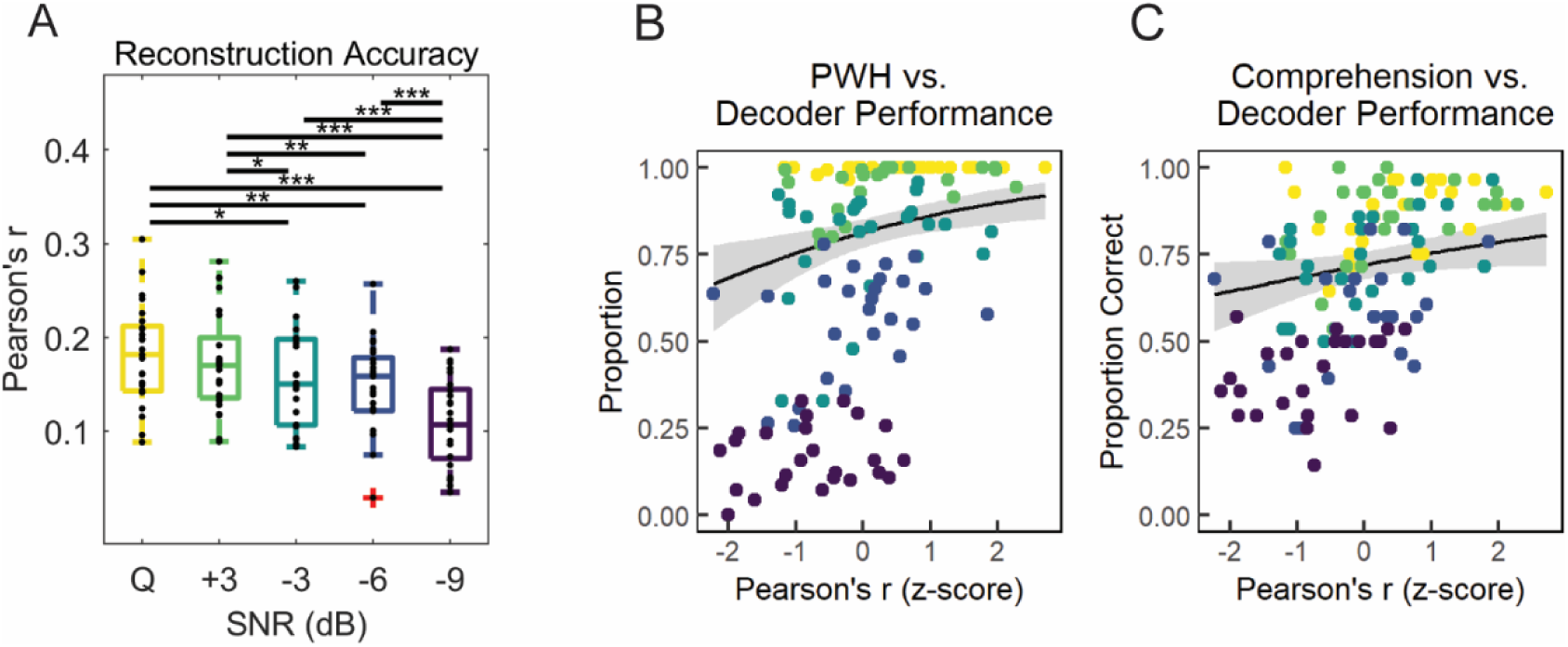
Envelope modeling results. (A) Colored boxplots show envelope reconstruction accuracies for each condition, and the black points represent individual participants. Significance is indicated by * if p < 0.05, ** if p < 0.01, and *** if p < 0.001 using pairwise t-tests with Holm correction. (B, C) The black lines indicate GLMM-predicted PWH (B) and comprehension scores (C) via *ggpredict* as a function of envelope tracking while controlling for SNR. The shaded regions represent the 95% confidence intervals, and each colored dot represents each participant’s average reconstruction accuracy in each SNR.

Next, we tested if the fidelity of envelope tracking was indicative of how well participants could hear and understand the story. This was tested using a GLMM model (Equation 5) that contained envelope reconstruction accuracies, SNR (using a linear polynomial contrast), and the interaction between the two as fixed effects and participants as the random effect. Envelope tracking showed a positive relationship with both PWH (β = 0.356, p = 1.991 x 10^-3^, R^2^_m_ = 0.926, R^2^_c_ = 0.996) and comprehension scores (β = 0.178, p = 0.040, R^2^_m_ = 0.804, R^2^_c_ = 0.983). Increases in SNR were also associated with increases in behavioral scores (PWH: β = 2.019, p = 2.268 x 10^-15^; comprehension: β = 0.840, p = 2.190 x 10^-6^). The interaction between reconstruction accuracy and SNR, however, was not significant in the PWH (β = 0.096, p = 0.247) GLMM nor the comprehension (β = 0.043, p = 0.416) GLMM. This indicates that the relationship between envelope tracking and behavioral scores does not vary across SNRs. In sum, neural tracking of the speech envelope provides a consistent predictor of both PWH and comprehension scores, independent of listening condition.

### Lexical surprisal’s influence on early auditory encoding decreases at high noise levels

Previous work by Broderick and colleagues has shown that the cortical tracking of an individual word’s acoustic representations was enhanced the more semantically similar it was to its preceding context (Broderick et al., 2019). Since higher-level representations can bias perception when incoming stimuli are noisy (de Lange et al., 2018), perhaps participants in the present study relied more on a higher-level feature (i.e., next word probability/surprisal) when noise levels slightly increased—thereby strengthening the influence of word surprisal on lower-level feature encoding. This influence of lexical surprisal should then decrease the noisier the speech becomes, i.e., as people begin to fail to understand the speech. We aimed to test these hypotheses using a two-stage regression analysis that consisted of stimulus envelope reconstruction followed by linear mixed effects modeling.

The backward modeling technique helps improve overall signal to noise ratio by simultaneously training on all 128 electrodes to reconstruct the (single channel) speech envelope. Because of this benefit, backward models were used in the two-stage regression rather than forward models. The decoders were used to reconstruct an estimate of the speech envelopes. Spearman’s correlation was then computed between the first 100 ms of the reconstructed and actual envelopes for each individual word. All words in the story were scored on their lexical surprisal, which was calculated as the negative logarithm of a word’s probability given the words that came before it. Envelope variability, relative pitch, and resolvability (the nuisance regressors) were also calculated to control for acoustic related properties of the stimuli, many of which could correlate with word surprisal and word reconstruction accuracy. Surprisal, SNR, and the nuisance regressors were included in an LME model to predict word reconstruction accuracy for the first 100 ms of each word.

There was a significant positive relationship between lexical surprisal and word reconstruction accuracy in quiet (β = 2.850 x 10^-2^, p < 2.22 x 10^-16^, R^2^_m_ = 0.017, R^2^ = 0.030, Table 2). In other words, the more surprising a word (or the less probable that word was given its preceding context), the better that word’s envelope was decoded from the EEG. The proportion of variance explained by the present LME model, R^2^_m_ = 0.017, is consistent with what was found by Broderick and colleagues, R^2^ = 0.0171 (Broderick et al., 2019). The interaction between surprisal and SNR shows how the *surp* coefficient changes (or is adjusted) across noise levels (Figure 6A, Table 2, S3 Table). For example, whereas surprisal in quiet has a coefficient of 2.850 x 10^-2^, surprisal in the +3 dB condition has a coefficient (or slope) that is 2.172x 10^-4^ units lower (S3 Table, top). The influence of lexical surprisal on word reconstruction accuracy in quiet was similar to the +3 dB (β = −2.172x 10^-4^, p = 0.951) and −3 dB (β = −4.767x 10^-3^, p = 0.173) SNR conditions and greater than the −6 dB (β = −9.900 x 10^-3^, p = 0.005) and −9 dB (β = −1.600 x 10^-2^, p = 4.890 x 10^-6^, Table 2) SNR conditions. This shows that lexical surprisal impacts envelope tracking in quiet and in low levels of background noise (+3 dB and −3 dB SNRs), but that this influence significantly decreases with moderate to high levels of noise. It is also important to note that the nuisance regressors and additional interactions included in the LME model also significantly influenced word reconstruction accuracy, reinforcing the value of controlling for those variables and ensuring our results were not confounded by other acoustic related measures.

**Figure 6.**
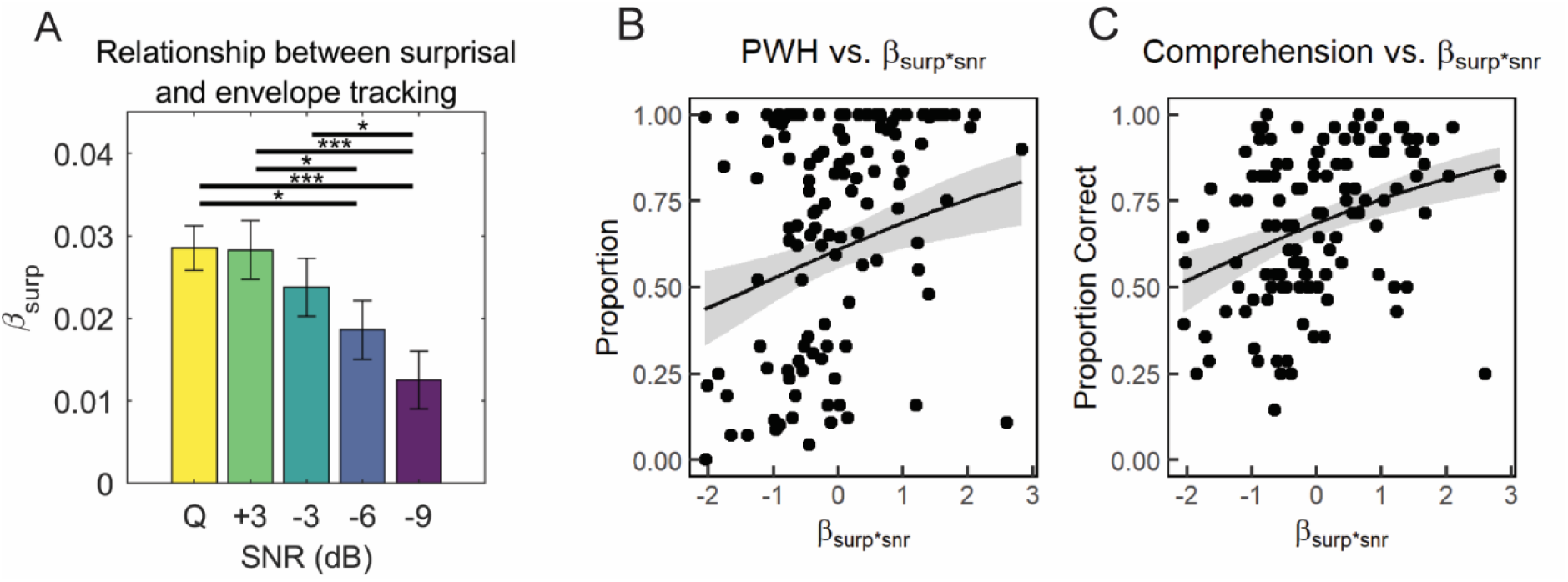
The relationship between lexical surprisal and envelope tracking at different levels of noise. (A) The surprisal coefficient in quiet and the interaction between surprisal and SNR using a 100 ms word window. The error bars are the standard errors of each measure, calculated using the LME model. Significance is indicated by * if p < 0.05, ** if p < 0.01, and *** if p < 0.001 using R’s *emtrends* function (*emmeans* package) to compare estimated marginal means of linear trends. (B, C) The relationship between the surprisal coefficients across SNRs and behavior. The percentage of words heard (PWH) scores are in B and comprehension scores are in C.

**Table 2.**
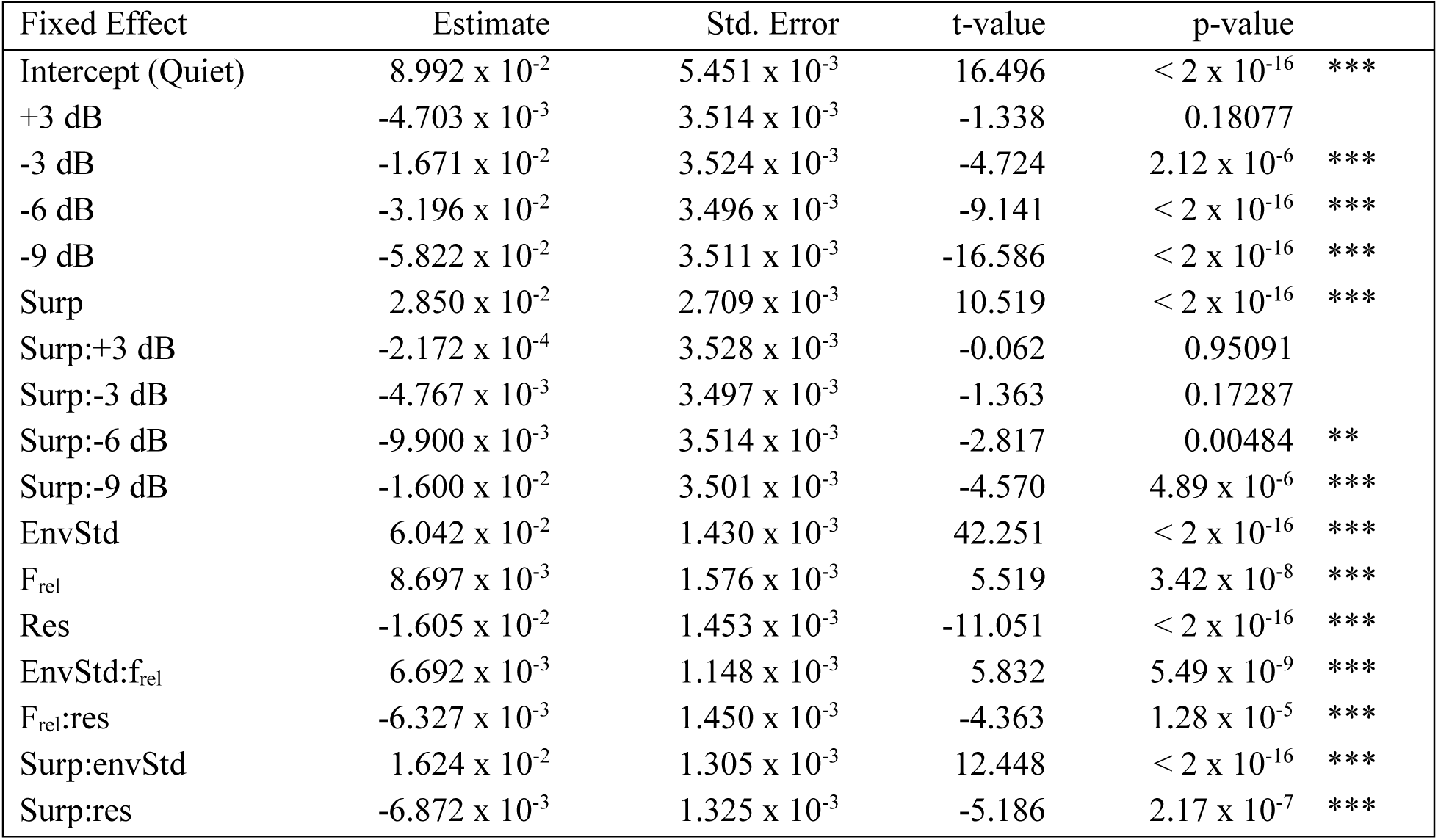
LME model showing the relationship between lexical surprisal and envelope tracking in each SNR.

| Fixed Effect | Estimate | Std. Error | t-value | p-value |  |
| --- | --- | --- | --- | --- | --- |
| Intercept (Quiet) | $8.992 \times 10^{-2}$ | $5.451 \times 10^{-3}$ | 16.496 | $< 2 \times 10^{-16}$ | *** |
| +3 dB | $-4.703 \times 10^{-3}$ | $3.514 \times 10^{-3}$ | -1.338 | 0.18077 | |
| -3 dB | $-1.671 \times 10^{-2}$ | $3.524 \times 10^{-3}$ | -4.724 | $2.12 \times 10^{-6}$ | *** |
| -6 dB | $-3.196 \times 10^{-2}$ | $3.496 \times 10^{-3}$ | -9.141 | $< 2 \times 10^{-16}$ | *** |
| -9 dB | $-5.822 \times 10^{-2}$ | $3.511 \times 10^{-3}$ | -16.586 | $< 2 \times 10^{-16}$ | *** |
| Surp | $2.850 \times 10^{-2}$ | $2.709 \times 10^{-3}$ | 10.519 | $< 2 \times 10^{-16}$ | *** |
| Surp:+3 dB | $-2.172 \times 10^{-4}$ | $3.528 \times 10^{-3}$ | -0.062 | 0.95091 | |
| Surp:-3 dB | $-4.767 \times 10^{-3}$ | $3.497 \times 10^{-3}$ | -1.363 | 0.17287 | |
| Surp:-6 dB | $-9.900 \times 10^{-3}$ | $3.514 \times 10^{-3}$ | -2.817 | 0.00484 | ** |
| Surp:-9 dB | $-1.600 \times 10^{-2}$ | $3.501 \times 10^{-3}$ | -4.570 | $4.89 \times 10^{-6}$ | *** |
| EnvStd | $6.042 \times 10^{-2}$ | $1.430 \times 10^{-3}$ | 42.251 | $< 2 \times 10^{-16}$ | *** |
| F<sub>rel</sub> | $8.697 \times 10^{-3}$ | $1.576 \times 10^{-3}$ | 5.519 | $3.42 \times 10^{-8}$ | *** |
| Res | $-1.605 \times 10^{-2}$ | $1.453 \times 10^{-3}$ | -11.051 | $< 2 \times 10^{-16}$ | *** |
| EnvStd:f<sub>rel</sub> | $6.692 \times 10^{-3}$ | $1.148 \times 10^{-3}$ | 5.832 | $5.49 \times 10^{-9}$ | *** |
| F<sub>rel</sub>:res | $-6.327 \times 10^{-3}$ | $1.450 \times 10^{-3}$ | -4.363 | $1.28 \times 10^{-5}$ | *** |
| Surp:envStd | $1.624 \times 10^{-2}$ | $1.305 \times 10^{-3}$ | 12.448 | $< 2 \times 10^{-16}$ | *** |
| Surp:res | $-6.872 \times 10^{-3}$ | $1.325 \times 10^{-3}$ | -5.186 | $2.17 \times 10^{-7}$ | *** |

Despite studies showing that low SNRs benefit from the utilization of context (Mayo et al., 1997), our finding that lexical surprisal significantly influences envelope tracking for the −6 dB (z-ratio = 6.814, p < 0.001) and −9 dB conditions (z-ratio = 4.615, p < 0.001, Holm corrected test that each SNR by surprisal trend is greater than 0, S3 Table) is unexpected. Participants reported hearing either no words or very few words in, for instance, the −9 dB SNR condition, yet we still see a significant influence of lexical surprisal on envelope tracking. We did not control for any other features (including surprisal) in our envelope reconstruction model, so these unaccounted-for features may have contributed to the significantly positive interaction coefficients in the −6 dB and −9 dB SNR conditions. Or the fact that some participants were able to hear some words in both conditions may have been enough to cause this significant effect. Lastly, we found that single subject surprisal coefficients also predicted their self-reported percentage of words heard (β = 0.358, p = 0.001, R^2^_m_ = 0.384, Figure 6B) and comprehension scores (β = 3.558, p = 1.906 x 10^-5^, R^2^_m_ =0.533, R^2^_c_ = 686) across SNRs (Equation 6, Figure 6C). That is to say, the stronger the influence of surprisal on envelope tracking, the better the participants were able to hear and comprehend the story.

## DISCUSSION

This study sought to establish how well indices of hierarchical neural speech processing reflect language comprehension—advancing on prior work that has typically tested specific hierarchical levels without controlling for the others. First, the encoding of a range of hierarchical speech features was characterized to assess how this encoding changed with noise and if those changes related to behavior. Linguistic feature representations were more affected by noise than the acoustic representations. In addition, associations between behavioral performance and speech feature tracking were task- and condition-dependent: PWH scores became decreasingly linked to acoustic onset tracking and increasingly linked to lexical surprisal tracking as SNR improved, whereas comprehension scores became increasingly linked to phonetic feature as SNR worsened. Speech envelope tracking fidelity was significantly associated with behavioral scores, and this relationship remained consistent across SNRs. The final set of analyses investigated the influence of lexical surprisal on neural tracking of the speech envelope. Results showed that the envelopes of more unexpected words were strongly reflected in the EEG in quiet and in low levels of noise, but this relationship weakened as noise levels increased.

### Neural representations of acoustic and phonetic features show dissociable sensitivity to noise

Acoustic feature representations were hypothesized to be more invariant to background noise than linguistic feature representations; this pattern was supported by the present results. The degree to which spectrogram information was reflected in EEG remained unchanged across SNRs. This result contrasts with studies such as Lesenfants and colleagues whose spectrogram model performance increased monotonically (as noise levels *decreased*) when training on delta band EEG and increased in a more S-shaped pattern when training on theta band EEG (Lesenfants et al., 2019). These differences may be partly explained by the efforts in the present study to partial out the contribution of other speech features when assessing model performance.

Interestingly, phonetic features were uniquely represented in neural activity even when accounting for the speech spectrogram and acoustic onsets. This is consistent with previous studies showing that the addition of phonetic features to spectrotemporal representations improves EEG prediction (Di Liberto et al., 2015; Sohoglu & Davis, 2020; Tezcan et al., 2023) and correlations with speech intelligibility (Lesenfants et al., 2019), and that phonetic features can uniquely predict EEG responses even when attending to a specific talker (Teoh et al., 2022). However, the present phonetic feature results contrast with previous work which suggested that responses to articulations could be explained by simpler acoustic features (Daube et al., 2019). Moreover, the differential decline of phonetic feature and spectrogram encoding, along with their distinct relationships to behavior, provides further evidence that the two features are dissociable.

When assessing speech envelope decoding on the other hand, the reconstruction accuracies first began to decrease in the −3 dB SNR and decreased yet again in the −9 dB SNR. This pattern was surprising given a prior report that the synchronization between neural activity and the speech envelope remained largely unaffected until the speech signal reached an SNR of −9 dB (Ding & Simon, 2013). These contrasting results may be attributed to a combination of factors: neural recording modality, data preprocessing, model training and testing procedures between conditions, or the regularization method used (e.g., boosting versus ridge regression). Instead, the present results show a gradual decrease in envelope tracking across SNRs similar to findings from Vanthornhout and colleagues (Vanthornhout et al., 2018). However, the present results indicate that envelope reconstruction accuracies were significantly associated with participants’ behavioral scores which is consistent with many studies that have shown that the speech envelope (using stimulus reconstruction or cross-correlation) contributes and relates to speech intelligibility and comprehension (Ahissar et al., 2001; Decruy et al., 2020; Iotzov & Parra, 2019; Lesenfants et al., 2019; Muncke et al., 2022; Vanthornhout et al., 2018).

### Behavioral relevance of acoustic and linguistic features varies with noise level

When further examining the relationship between feature-specific neural tracking and behavioral performance, the present results revealed a more nuanced pattern than originally hypothesized. Rather than observing a uniform dominance of higher-level linguistic feature tracking across listening conditions, the behavioral relevance of neural representations depended on both listening condition and the behavioral metric being assessed. The association between what participants reported hearing and lexical surprisal tracking strengthened as speech became cleaner (less noisy). This finding is consistent with prior work which demonstrated that lexical surprisal tracking is prominent when speech is comprehensible (Gillis et al., 2023; Mesik et al., 2021) and that this tracking weakens as speech becomes harder to understand (Verschueren et al., 2022). In contrast, the relationship between PWH and acoustic onset tracking weakened as speech became cleaner. This suggests that this low-level cue was more behaviorally relevant when the speech signal was degraded (Davis & Johnsrude, 2007; Mattys et al., 2005).

A different pattern was observed for comprehension scores. Both spectrogram and lexical surprisal based representations were positively associated with comprehension scores across listening conditions, suggesting stable contributions of acoustic and linguistic processing to speech understanding. Notably, however, only phonetic feature tracking showed a significant interaction with SNR such that its association with comprehension decreased as noise levels decreased. This pattern indicates that the ability to convert acoustic speech sounds into phonetic-level representations in noisy conditions supports comprehension of that speech under those conditions. Across behavioral measures, these findings suggest that the behavioral relevance of neural representations shifts across hierarchical levels as listening conditions change. Specifically, lower-level acoustic and phonetic information contributes more strongly under noisy conditions, whereas higher-level linguistic predictions become most informative when speech is in quiet. Consistent with prior work showing that speech in noise perception is best predicted by models that combine neural measures across multiple levels (Lesenfants et al., 2019), the present study complements this finding by showing that speech perception and comprehension rely in parallel on acoustic and phonetic representations. Our approach reveals how the behavioral relevance of hierarchical speech feature tracking is differentially weighted across listening conditions, an insight that may be less apparent in models combining multiple speech features.

### Lexical context can predict the fidelity of word-level acoustic tracking, but this effect decreases as noise levels increase

Another key hypothesis of our study was that participants would use lexical context to predict and encode the acoustic features of each word. This hypothesis was supported: the LME analysis (stage two of the two-stage regression) revealed that the more unexpected a word, the better that word’s envelope was reconstructed. However, we also hypothesized that participants would rely more on these predictions for speech in moderate levels of noise (when speech is still intelligible) relative to speech in quiet, before declining at high levels of background noise (when speech is no longer intelligible). This result was only partially borne out. Specifically, the use of lexical context in processing speech acoustics did decrease as the speech became noisier, but there was no evidence to support a stronger reliance on context in moderate levels of noise. In particular, while there was no difference in comprehension scores between the quiet and +3 dB conditions, there was no increase in the influence of surprisal on envelope tracking for the latter condition compared to speech in quiet. However, a larger spread of the percentage of words heard scores was observed across participants in the +3 dB condition. This raised the possibility that participants who were starting to struggle might put forth more effort to understand and process the story by relying more on context (and thus might have a higher surprisal weight in Figure 6A) than those who remained at ceiling. Nevertheless, no significant difference was observed to support this possibility. This was a little surprising given that context has been known to affect behavior (Golestani et al., 2013) and neural activity (Boulenger et al., 2011; Koskinen et al., 2020; Strauß et al., 2022) in challenging listening conditions. Future work with larger participant numbers and perhaps even lower levels of background noise (e.g., + 6 dB SNR) might reveal such an effect.

The present second-stage regression analysis based on word surprisal seems to be at odds with Broderick and colleagues who found that the envelopes of words that were more semantically *similar* to their context were better reflected in the EEG. That is to say, envelope tracking is enhanced for words that share a similar meaning with their context (Broderick et al., 2019). However, semantic similarity and lexical surprisal tend to share a moderate, and sometimes weak negative correlation (Frank & Willems, 2017). Indeed, a re-analysis of Broderick et al.’s original EEG data has revealed that both semantic similarity and lexical surprisal play complementary (positive) roles in estimating when envelope tracking is enhanced (Broderick & Lalor, 2020). Nevertheless, the nature of this duality remains mysterious, and we hope it will provide the grounds for an exciting body of future work. We anticipate it will take a substantial battery of future experiments to shape a unifying explanation, with stimuli that can disentangle correlations between semantic similarity, lexical surprisal, and other linguistic factors that could come into play (e.g., semantic content, next-word entropy, phonetic surprisal, next-phoneme entropy). In any case, what seems clear in the present results is that lexical context influences the neural tracking of speech acoustics on a word-by-word basis, and this influence drops as speech becomes unintelligible.

### Methodological limitations

The present study aimed to characterize how hierarchical speech feature encoding changes in increasing levels of background noise. One inherent challenge was the selection of speech features, particularly given the relatively low signal to noise ratio of EEG and the limited amount of EEG data per participant. Numerous candidate features have been used in prior work such as the spectrogram derivative (Teoh et al., 2022), log-mel spectrogram (Daube et al., 2019), phoneme onset, cohort entropy (Brodbeck, Hong, et al., 2018), and semantic dissimilarity (Broderick et al., 2018). As such, we opted to focus on a set of commonly used features spanning acoustic, phonetic, and lexical levels to balance interpretability with model complexity.

A related challenge is that speech features are likely to correlate with one another and explain similar activity in the EEG. This motivated our use of the partial correlation analyses to estimate the unique contribution of each feature to the neural responses while controlling for the effect of others. While this approach provides a principled way to disentangle correlated representations, results will partly depend on the specific feature set included.

Another potential limitation of this study was the mode of regularization. We optimized the regularization parameter on the clean speech and used this same parameter on all other conditions. This choice was motivated by the desire to reduce overfitting, as this would be particularly prudent at lower SNRs (i.e., −6 dB and −9 dB) where multidimensional feature models would be more susceptible to noise-driven fits. Although this approach may reduce sensitivity to condition-specific changes in optimal regularization, it provides a conservative and unbiased basis for comparing model performance across listening conditions.

## Conclusion

In summary, the current results showed that neural representations of lexical surprisal and phonetic features were more susceptible to noise than acoustic speech feature representations. Behavioral performance reflected contributions from multiple hierarchical levels of speech processing, with the relative predictive value of acoustic, phonetic, and lexical features varying with noise level and task. We also found that the encoding of certain phonetic features decreased even in low levels of noise, and that the encoding of frequencies below 1.3k essentially disappeared in high levels of noise. Lastly, we showed that context—in the form of a word predictability—influenced how the acoustics of those words were encoded. This influence lessens in high background noise levels. Future work will aim to further characterize how people might rely more or less on top-down context to process bottom-up speech input as a function of stimulus type, task, and listening conditions.

Author Contributions

ECL and SRS designed research; SRS performed research; AJA contributed analytic tools, SRS analyzed data; SRS, AJA, ECL wrote the paper

## Acknowledgements

The authors thank Ms. Xueying Wang and Dr. Aaron Nidiffer for some assistance with data collection, and Dr. Aaron Nidiffer and Dr. Madeline Cappelloni for helpful comments on the manuscript.

Authors report no conflict of interest.

## SUPPLEMENTARY INFORMATION

**S1 Table.**
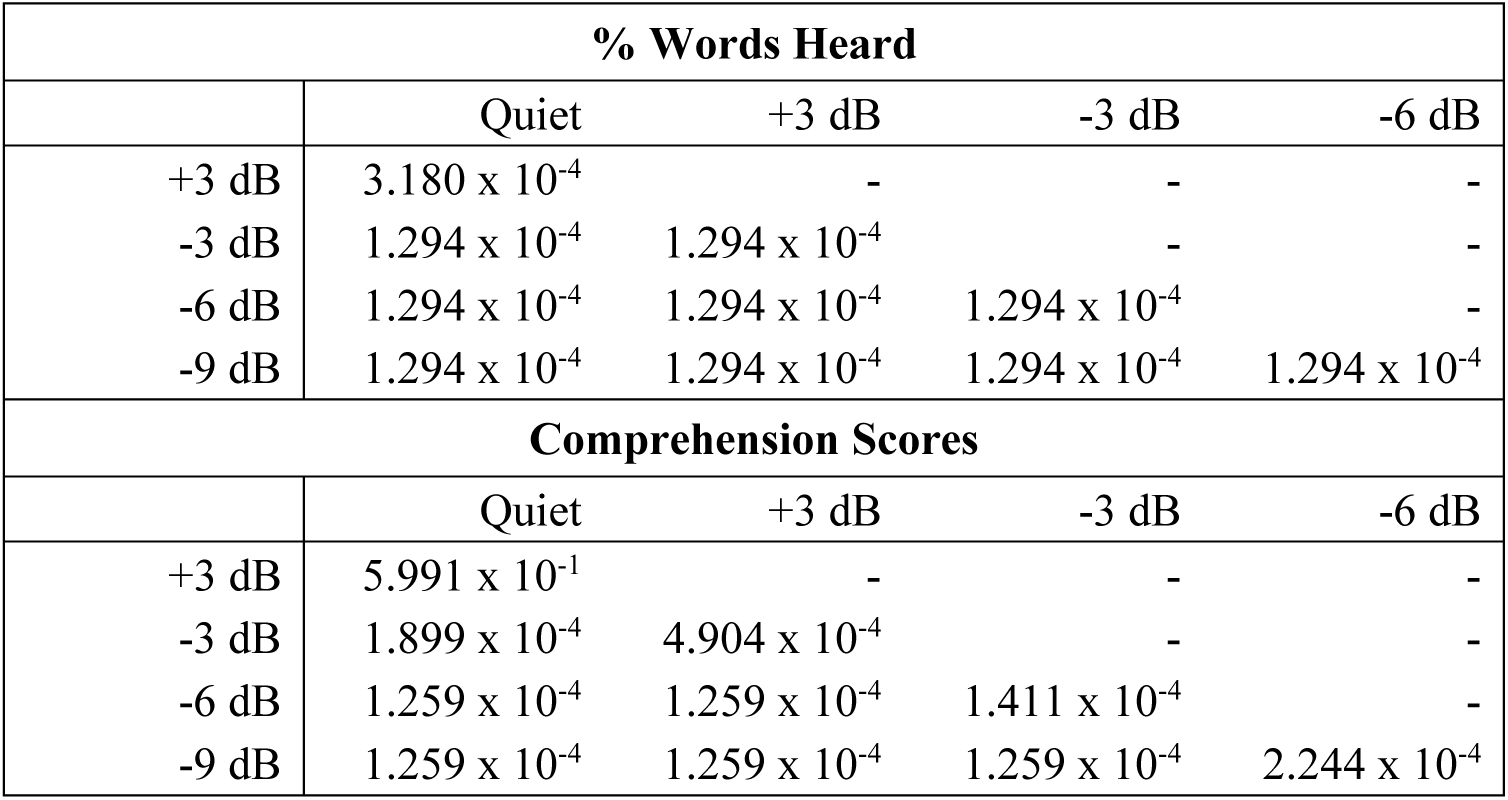
Wilcoxon rank sum p-values from behavioral score comparisons between conditions.

| % Words Heard |  |  |  |  |
| --- | --- | --- | --- | --- |
|  | Quiet | +3 dB | -3 dB | -6 dB |
| +3 dB | $3.180 \times 10^{-4}$ | - | - | - |
| -3 dB | $1.294 \times 10^{-4}$ | $1.294 \times 10^{-4}$ | - | - |
| -6 dB | $1.294 \times 10^{-4}$ | $1.294 \times 10^{-4}$ | $1.294 \times 10^{-4}$ | - |
| -9 dB | $1.294 \times 10^{-4}$ | $1.294 \times 10^{-4}$ | $1.294 \times 10^{-4}$ | $1.294 \times 10^{-4}$ |
| Comprehension Scores |  |  |  |  |
|  | Quiet | +3 dB | -3 dB | -6 dB |
| +3 dB | $5.991 \times 10^{-1}$ | - | - | - |
| -3 dB | $1.899 \times 10^{-4}$ | $4.904 \times 10^{-4}$ | - | - |
| -6 dB | $1.259 \times 10^{-4}$ | $1.259 \times 10^{-4}$ | $1.411 \times 10^{-4}$ | - |
| -9 dB | $1.259 \times 10^{-4}$ | $1.259 \times 10^{-4}$ | $1.259 \times 10^{-4}$ | $2.244 \times 10^{-4}$ |

**S1 Figure.**
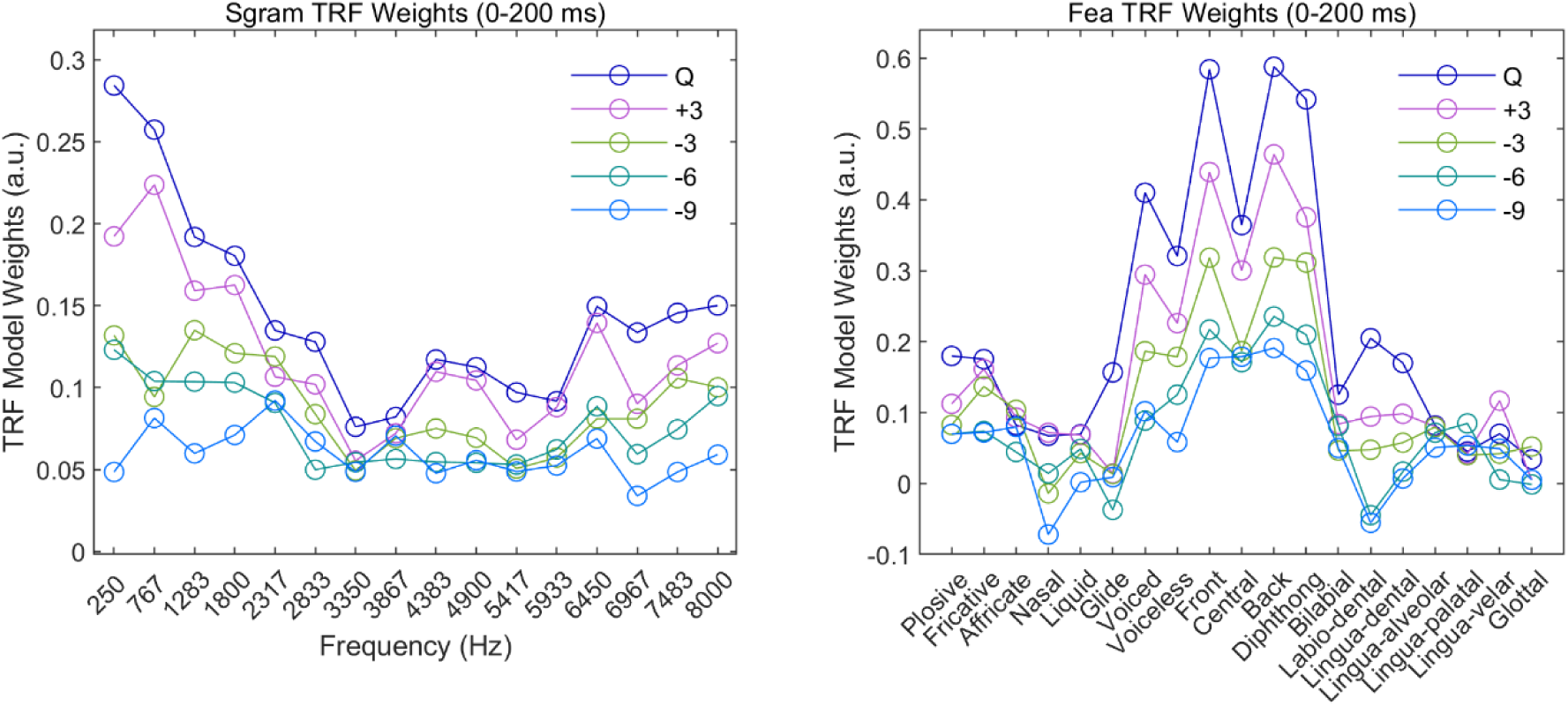
Spectrogram (Sgram, left) and Phonetic Feature (Fea, right) TRF model weights at each signal to noise ratio. The model weights were averaged across time (0 to 200 ms) and the 12 frontotemporal electrodes depicted in Figure 1. Frequency bands and feature spaces are plotted on the x-axes and model weights (in arbitrary units) are on the y-axes.

**S2 Table.** Results from GLMMs that related partial correlation coefficients to behavioral scores.

| Percent words heard score GLMM |  |  |  |  |  |
| --- | --- | --- | --- | --- | --- |
| Fixed Effect | Estimate | Std. Error | t-value | p-value |  |
| Intercept | 1. 609 | 0.138 | 11.635 | < 2 x 10<sup>-16</sup> | *** |
| Ons<sub>A</sub> | -0.129 | 0.086 | -1.497 | 0.134 |  |
| Sgram | 0.117 | 0.088 | 1.322 | 0.186 |  |
| Fea | 0.104 | 0.102 | 1.022 | 0.307 |  |
| Surp | 0.398 | 0.097 | 4.128 | 3.652 x 10<sup>-5</sup> | *** |
| Ons<sub>W</sub> | 0.055 | 0.070 | 0.779 | 0.436 |  |
| SNR | -2.133 | 0.104 | -20.454 | < 2 x 10<sup>-16</sup> | *** |
| SNR:Ons<sub>A</sub> | 0.258 | 0.092 | 2.819 | 0.005 | ** |
| SNR:Sgram | -0.038 | 0.073 | -0.528 | 0.597 |  |
| SNR:Fea | -0.098 | 0.111 | -0.877 | 0.381 |  |
| SNR:Surp | -0.265 | 0.098 | -2.694 | 0.007 | ** |
| SNR:Ons<sub>W</sub> | -0.083 | 0.071 | -1.165 | 0.243 |  |
| Comprehension score GLMM |  |  |  |  |  |
| Fixed Effect | Estimate | Std. Error | t-value | p-value |  |
| Intercept | 1.046 | 0.111 | 9.402 | < 2 x 10<sup>-16</sup> | *** |
| Ons<sub>A</sub> | -0.015 | 0.062 | -0.244 | 0.807 |  |
| Sgram | 0.128 | 0.063 | 2.031 | 0.042 | * |
| Fea | 0.055 | 0.060 | 0.919 | 0.358 |  |
| Surp | 0.122 | 0.061 | 1.996 | 0.046 | * |
| Ons<sub>W</sub> | 0.044 | 0.051 | 0.871 | 0.384 |  |
| SNR | -0.830 | 0.070 | -11.932 | < 2 x 10<sup>-16</sup> | *** |
| SNR:Ons<sub>A</sub> | 0.033 | 0.055 | 0.598 | 0.550 |  |
| SNR:Sgram | -0.051 | 0.049 | -1.022 | 0.307 |  |
| SNR:Fea | 0.124 | 0.058 | 2.137 | 0.033 | * |
| SNR:Surp | 0.024 | 0.053 | 0.462 | 0.644 |  |
| SNR:Ons<sub>W</sub> | 0.038 | 0.050 | -0.777 | 0.437 |  |

**S3 Table.** Post hoc contrasts for the LME model that calculated the relationship between lexical surprisal and envelope tracking in each SNR.

| Interaction variable statistics |  |  |  |  |  |
| --- | --- | --- | --- | --- | --- |
| SNR | Surp:SNR Slope | Std. Error | z-ratio | p-value |  |
| Quiet | 0.029 | 2.709 x 10<sup>-3</sup> | 10.519 | < 0.001 | *** |
| +3 dB | 0.028 | 2.701 x 10<sup>-3</sup> | 10.336 | < 0.001 | *** |
| -3 dB | 0.024 | 2.730 x 10<sup>-3</sup> | 8.787 | < 0.001 | *** |
| -6 dB | 0.019 | 2.709 x 10<sup>-3</sup> | 6.814 | < 0.001 | *** |
| -9 dB | 0.013 | 2.736 x 10<sup>-3</sup> | 4.615 | < 0.001 | *** |
| Comparisons between interaction variables |  |  |  |  |  |
| Contrast | Estimate | Std. Error | z ratio | p-value |  |
| Quiet – + 3 dB | 2.17 x 10<sup>-4</sup> | 3.53 x 10<sup>-3</sup> | 0.062 | 1.000 |  |
| Quiet – -3 dB | 4.767 x 10<sup>-3</sup> | 3.50 x 10<sup>-3</sup> | 1.363 | 0.651 |  |
| Quiet – -6 dB | 9.900 x 10<sup>-3</sup> | 3.51 x 10<sup>-3</sup> | 2.817 | 0.039 | * |
| Quiet – -9 dB | 1.600 x 10<sup>-2</sup> | 3.50 x 10<sup>-3</sup> | 4.570 | < 0.001 | *** |
| -3 dB – + 3 dB | -4.459 x 10<sup>-3</sup> | 3.53 x 10<sup>-3</sup> | -1.290 | 0.698 |  |
| -3 dB – -6 dB | 5.133 x 10<sup>-3</sup> | 3.52 x 10<sup>-3</sup> | 1.458 | 0.590 |  |
| -3 dB – -9 dB | 1.123 x 10<sup>-2</sup> | 3.50 x 10<sup>-3</sup> | 3.211 | 0.012 | * |
| -6 dB – + 3 dB | -9.683 x 10<sup>-3</sup> | 3.53 x 10<sup>-3</sup> | -2.732 | 0.049 | * |
| -6 dB – -9 dB | 6.097 x 10<sup>-3</sup> | 3.54 x 10<sup>-3</sup> | 1.729 | 0.416 |  |
| -9 dB – + 3 dB | -1.578 x 10<sup>-2</sup> | 3.53 x 10<sup>-3</sup> | -4.468 | < 0.001 | *** |

